# The Arp2/3 complex and the formin, Diaphanous, are both required to regulate the size of germline ring canals in the developing egg chamber

**DOI:** 10.1101/813873

**Authors:** Josephine Thestrup, Marina Tipold, Alexandra Kindred, Kara Stark, Travis Curry, Lindsay Lewellyn

## Abstract

Intercellular bridges are an essential structural feature found in both germline and somatic cells throughout the animal kingdom. Because of their large size, the germline intercellular bridges, or ring canals, in the developing fruit fly egg chamber are an excellent model to study the formation, stabilization, and expansion of these structures. Within the egg chamber, the germline ring canals connect 15 supporting nurse cells to the developing oocyte, facilitating the transfer of materials required for successful oogenesis. The ring canals are derived from a stalled actomyosin contractile ring; once formed, additional actin and actin-binding proteins are recruited to the ring to support the 20-fold expansion that accompanies oogenesis. These behaviors provide a unique model system to study the actin regulators that control incomplete cytokinesis, intercellular bridge formation, and expansion. By temporally controlling their expression in the germline, we have demonstrated that the Arp2/3 complex and the formin, Diaphanous (Dia), coordinately regulate ring canal size and expansion throughout oogenesis. Dia is required for successful incomplete cytokinesis and the initial stabilization of the germline ring canals. Once the ring canals have formed, the Arp2/3 complex and Dia cooperate to determine ring canal size and maintain their stability. Our data suggest that the nurse cells must maintain a precise balance between the activity of these two nucleators during oogenesis.

## Introduction

Intercellular bridges are structures that connect neighboring cells, facilitating the transfer of materials and coordination of behaviors. These structures are found throughout the animal kingdom in both somatic and germ cells, and in many animals, their presence is essential for gametogenesis. In general, intercellular bridges are quite small, typically 0.5 µm to 2 µm in diameter. However, the germline intercellular bridges in the developing fruit fly egg chamber can be as large as 10 µm (Fawcett et al., 1959; Greenbaum et al., 2011; Haglund et al., 2011; Robinson and Cooley, 1996; Spradling, 1993), which makes them an ideal model system to study the mechanisms that contribute to formation, stabilization, and growth.

Each mature fruit fly egg arises from a structure called an egg chamber. Egg chamber formation begins in the germarium with the asymmetric division of a germline stem cell. One of the daughters of the germline stem cell, the cystoblast cell, undergoes four rounds of mitosis to generate the 16-cell cyst. The cyst is encapsulated by somatic follicle cells, and buds from the germarium, passing through 14 distinct stages to form the mature egg. The mitotic divisions of the germ cells end with incomplete cytokinesis, whereby the stalled contractile ring is stabilized to build an intercellular bridge or ring canal. The ring canal undergoes both thickening and expansion throughout oogenesis in preparation for the process of nurse cell dumping in stage 11. During nurse cell dumping, actomyosin-based contractility rapidly transfers cytoplasmic material from the nurse cells to the oocyte (Ferreira et al., 2014; Gutzeit, 1986; Spradling, 1993; Wheatley et al., 1995). To withstand the force of this dumping, ring canal stabilization and expansion are essential; mutations that disrupt either aspect lead to incomplete cytoplasmic transfer and the formation of smaller, non-viable eggs.

Actin is an essential component of the germline ring canals, whose levels and structural organization change as oogenesis proceeds. Electron microscopy analysis has demonstrated that the actin filaments within the ring canals tend to be long (>3 µm in diameter) and packed parallel to each other at a constant density throughout oogenesis (Fig. 1A). Despite the relatively constant density, there are clear developmentally-controlled changes in the rate of actin filament nucleation and organization within the ring canal. During early oogenesis (Phase I), the filament number increases from ~80 at stage 2 to over 700 at stage 6; this increase in filament number is accompanied by a >6-fold increase in the thickness of the ring, but only a modest increase in diameter. Beginning in stage 6 (Phase II), the diameter increases more rapidly, but the number of actin filaments per ring canal remains fairly constant until nurse cell dumping (Fig. 1A; Tilney, 1996). During Phase II, actin within the ring canal turns over at a rate that is similar to that observed at the leading edge of the lamellipodia of migrating cells (Kelso et al., 2002; McGrath et al., 1998; Theriot and Mitchison, 1991). These observations suggest that the activity of one or more actin nucleators is regulating the stage-specific changes in actin filament nucleation and elongation during oogenesis.

**Fig. 1:**
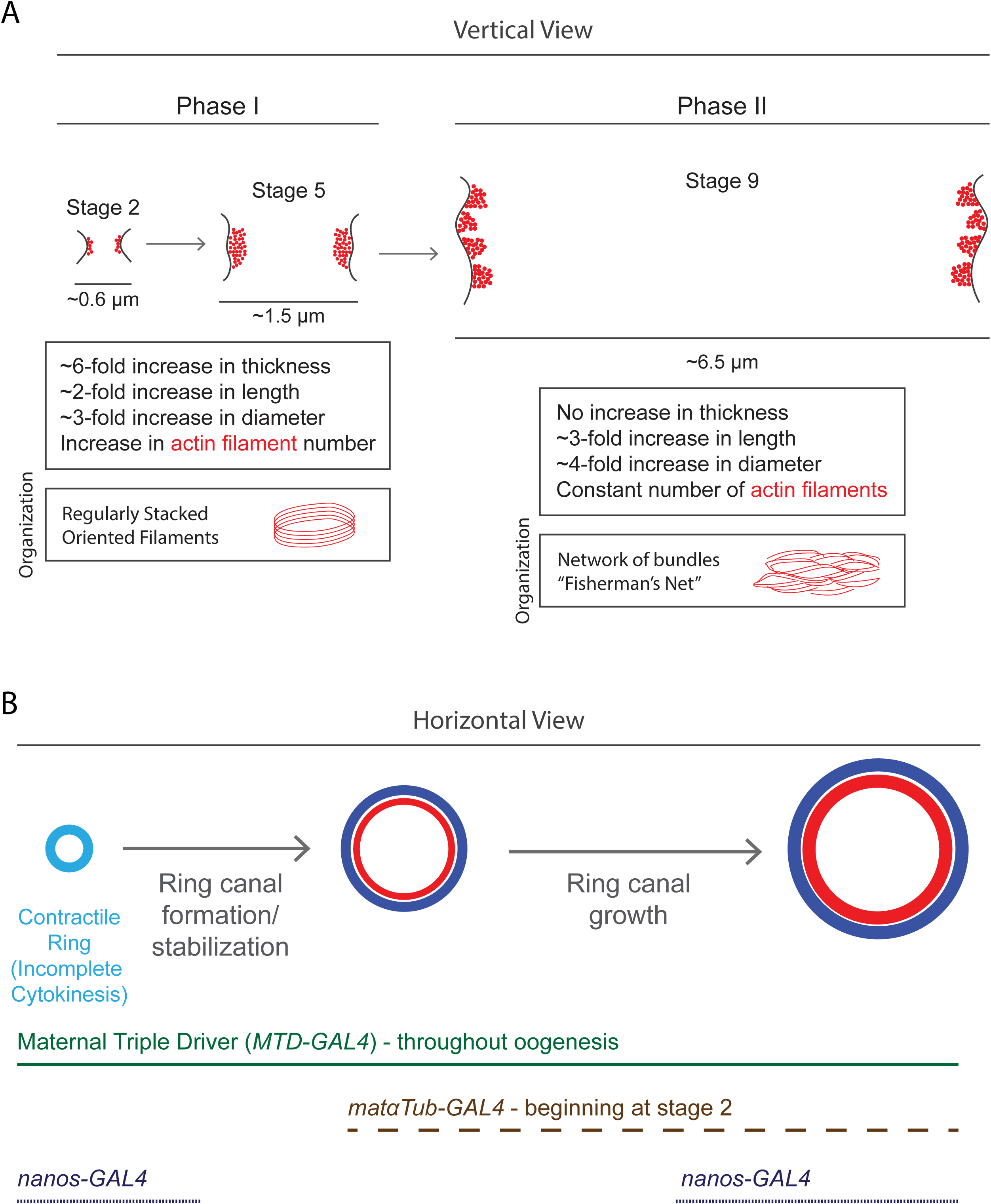
Stages of ring canal expansion and GAL4 drivers that can be used to regulate transgene expression. (A) Cartoon illustrating the structural changes that occur during the two phases (Phase I and Phase II) of ring canal expansion. Evidence for these two phases is based on electron microscopy (Tilney, 1996). (B) The temporal expression of GAL4 in each of the germline “driver” lines used in the study is indicated.

In addition to the stage-specific differences in actin filament number, the organization of the f-actin within the ring changes. During Phase I, the actin filaments are homogenously and tightly packed together. In contrast, during Phase II, the regular network of filaments is interrupted by spaces, evoking a fisherman’s net (Fig. 1A; Riparbelli and Callaini, 1995; Tilney, 1996). This reorganization suggests that there could be stage-specific differences in the activity or abundance of one or more actin bundlers, in addition to differences in the activity of one or more actin nucleators.

Despite this high-resolution analysis of f-actin organization and abundance within the ring canals, the proteins that nucleate and organize actin filaments have not been fully characterized. Most cells have two main types of actin nucleators. The Arp2/3 complex facilitates the formation of branched networks, which are essential for formation of lamellipodia during cell migration, for endocytosis, and recently have been implicated in cytokinesis (Chan et al., 2019; Pollard, 2007). In contrast, the formin family of actin nucleators can nucleate actin filaments, as well as promote their elongation by binding processively to the barbed end. Formin family members have been implicated in many morphogenetic processes and essential structures, such as in building the contractile ring during cytokinesis and in the formation of filopodia and lamellipodia in migrating cells (Breitsprecher and Goode, 2013; Pollard, 2007). The nature of the ring canals, which are derived from a stalled contractile ring that must undergo a ~20-fold increase in diameter, provides a unique model to study the role of different actin nucleators in multiple actin-based processes within a single cell type.

The Arp2/3 complex has been implicated in regulating ring canal size during oogenesis. In nurse cells homozygous for mutations in the Arp2/3 complex subunits ArpC1 and Arp3, ring canals are able to form but develop defects in size and shape, which were first visible around stage 5, which is at the end of Phase I (Hudson and Cooley, 2002). Similarly, mutation of the Arp2/3 activator, SCAR, was associated with smaller and collapsed ring canals (Zallen et al., 2002). These data suggest that SCAR-mediated activation of the Arp2/3 is required for ring canal expansion and stability during the final part of Phase I and during Phase II. However, the actin nucleator(s) that promote ring canal formation and expansion during the earlier part of Phase I have not been identified, nor is it known whether the Arp2/3 complex is the only nucleator required during Phase II. It is conceivable that a single actin nucleator, such as the Arp2/3 complex, could regulate ring canal expansion, or that the sequential or cooperative activity of multiple nucleators is required to regulate the developmentally controlled changes to the ring structure.

Here, we provide evidence that the Arp2/3 complex and the formin Diaphanous (Dia) play distinct roles in regulating the size and stability of the germline ring canals in the developing fruit fly egg. By using different GAL4s to temporally control transgene expression in the germline, we show that Dia is required for successful incomplete cytokinesis and initial formation of the germline ring canals. Once the ring canals have formed, the Arp2/3 complex and Dia cooperate to determine ring canal size and maintain stability, to facilitate efficient cytoplasmic transfer from the nurse cells to the oocyte.

## Results

### Depletion of ArpC2 leads to changes in ring canal size throughout oogenesis

Previous studies had demonstrated that germline mutations to the Arp2/3 complex subunits (*ArpC1* and *Arp3*) lead to smaller and collapsed ring canals, which were first observed at stage 5 of oogenesis (Hudson and Cooley, 2002). Based on this phenotype, it is possible that the Arp2/3 complex is dispensable for incomplete cytokinesis and the earlier part of Phase I; alternatively, the Arp2/3 complex may play a more subtle role in these earlier processes, which only becomes obvious by the end of Phase I.

In order to determine whether the Arp2/3 complex plays an earlier role in ring canal formation and/or Phase I expansion, we performed a strong depletion of the Arp2/3 complex subunit, ArpC2, throughout oogenesis using the Maternal Triple Driver (Fig. 1B; *MTD-GAL4*) and measured ring canal diameter from the germarium stage onwards. Ring canal diameter in the *arpC2-RNAi* egg chambers was notably variable compared to controls. Within the germarium, and through the first phase of ring canal expansion, the average diameter of ring canals was significantly higher in egg chambers depleted of ArpC2 (Fig. 2A-C). Beginning in stage 4, extremely small, lumenless ring canals also appeared (Fig. 2F). These “lumenless ring canals” were marked with phalloidin and Hts-RC but lacked a clear opening or lumen that would support the transfer of cytoplasmic materials from the nurse cells to the oocyte. Excluding lumenless ring canals, average ring canal diameter varied significantly from controls during most of Phase II (Fig. 2A-C). The mature eggs that developed from the ArpC2-depleted egg chambers were significantly smaller in volume than controls (Fig. 2G). These data suggest that the Arp2/3 complex may be necessary to modulate ring canal size throughout oogenesis, possibly beginning with an early, previously unappreciated role during incomplete cytokinesis of the cystoblast cells or during the earlier part of Phase I of ring canal growth.

**Fig. 2:**
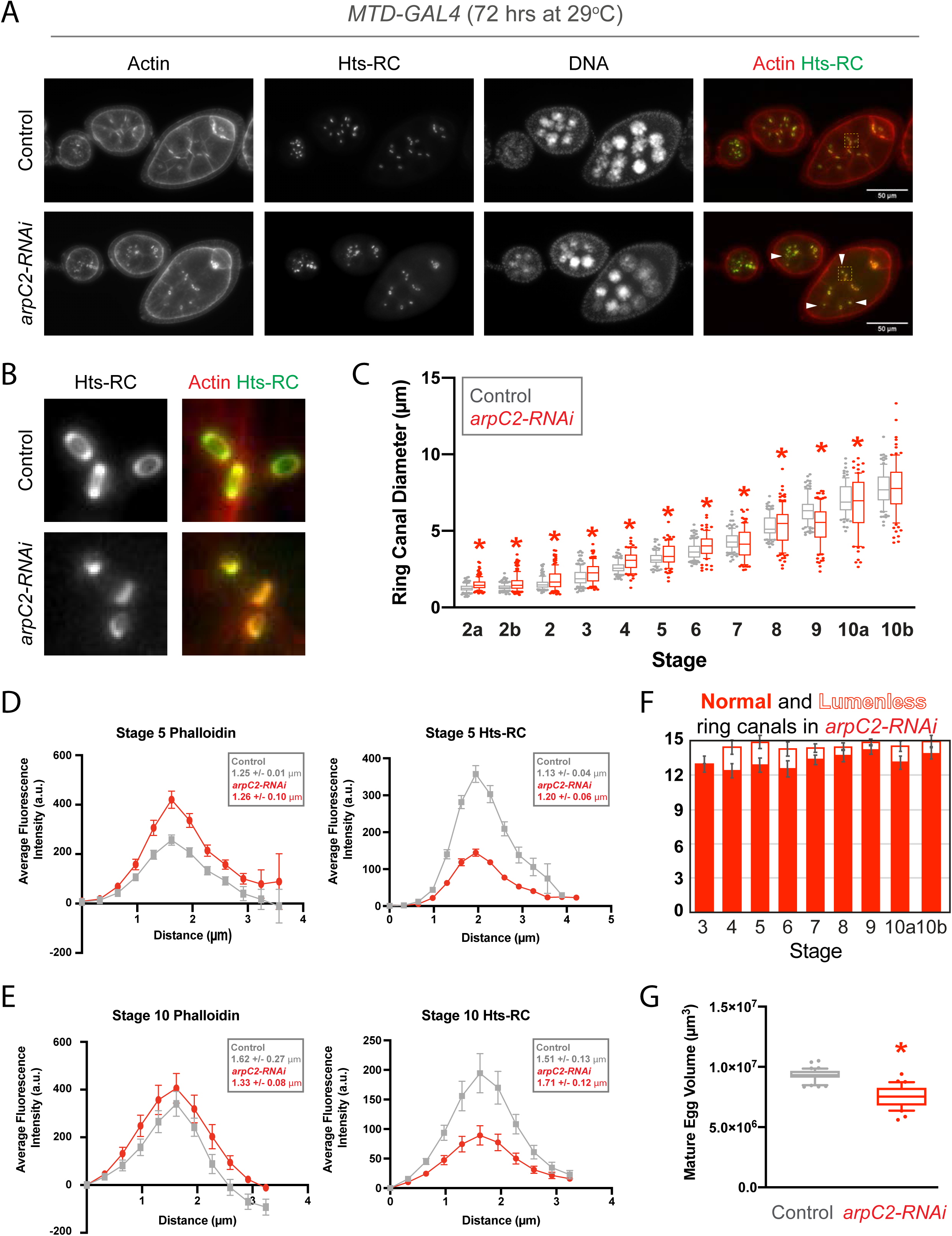
The Arp2/3 complex is required to regulate ring canal size throughout oogenesis. (A) Maximum intensity projections of control and *arpC2-RNAi* egg chambers. (B) Magnified view of the ring canals indicated by the boxed regions in A. (C) Box and whiskers plot showing the 10-90^th^ percent values for the diameter of ring canals connecting nurse cells. Individual points represent values outside of that range; lumenless ring canals were excluded from this analysis. n = 82-149 ring canals/stage for each condition. Asterisks indicate significant difference compared to control (p<0.05, K-S test). (D,E) Average fluorescence intensity of phalloidin and Hts-RC staining in ring canals of (D) stage 5 and (E) stage 10 control and *arpC2-RNAi* egg chambers (n=31 for control, n=39 for *arpC2-RNAi* in D; n=16 for control, n=17 for *arpC2-RNAi* in E). Error bars are SEM. Average full width at half maximum +/− SEM is shown for each stain in each condition. (F) Average number of total ring canals and lumenless ring canals in the *arpC2-RNAi* egg chambers at each stage. n=8-12 egg chambers per stage per condition. Error bars represent SEM. (G) Box and whiskers plot showing the 10-90^th^ percent values for the volumes of mature eggs. n=44 eggs for control and n=38 eggs for *arpC2-RNAi*. Asterisk indicates significant difference compared to control (p<0.0001, 2-tailed t-test). For all experiments, ArpC2 was depleted throughout oogenesis using the maternal triple driver (MTD-GAL4).

In order to determine whether depletion of ArpC2 affects the levels of actin and/or the ring canal component, Hts-RC, we performed linescan analysis on ring canals from control and *arpC2-RNAi* egg chambers. Interestingly, the ring canals from stage 5 *arpC2-RNAi* egg chambers contained higher levels of phalloidin within the ring canals, yet lower levels of the actin-binding protein, Hts-RC (Fig. 2D). During this analysis, we did observe a reduction in the cortical phalloidin signal in the stage 5 *arpC2-RNAi* condition, which could partially explain the relative increase in the ring canal signal. Thickness of the ring canals was approximated by averaging the full width at half maximum for each individual linescan; ring thickness did not differ significantly at this stage. At stage 10, Hts-RC within the ring was reduced, but levels of phalloidin and ring thickness were unchanged (Fig. 2E). Thus, the dramatic changes in the ring canal size in *arpC2-RNAi* egg chambers are not accompanied by severe defects in actin accumulation. It may be that another nucleator in the germline promotes actin nucleation during early stages to partially compensate for the loss of the Arp2/3 complex.

### Depletion of Dia leads to reduced ring canal number and changes in ring canal size

Although depletion of the Arp2/3 complex subunit, ArpC2, led to significant variability in ring canal size, the ring canals contained a normal amount of f-actin, and most egg chambers contained the expected number of ring canals (15 ring canals; 11 connecting nurse cells and 4 connecting the nurse cells to the oocyte). Therefore, we wondered whether another actin nucleator is required for incomplete cytokinesis and ring canal formation. Because of its role in cytokinesis and cellularization in the embryo (Afshar et al., 2000; Castrillon and Wasserman, 1994), we tested whether the formin Diaphanous (Dia) is required in the germline. Strong depletion of Dia using *MTD-GAL4* led to defects in ovary formation (*data not shown*); therefore, we utilized the *nanos-GAL4* driver under the control of the GAL80^ts^ inhibitor to suppress expression of the *UAS-dia-RNAi* transgene until adulthood; *nanos-GAL4* produces a pulse of transgene expression in the germline stem cells and then another pulse in mid-oogenesis (Fig. 1B; Hudson and Cooley, 2014). Egg chambers thus depleted of Dia in the germline often contained fewer than 15 ring canals, and these ring canals were significantly larger than in controls (Fig. 3A-C,F). In contrast to the ArpC2-depleted egg chambers, the ring canals from *dia-RNAi* egg chambers were reduced for both actin and Hts-RC at stage 5 (Fig. 3D) and contained slightly higher levels of actin at stage 10 (Fig. 3E). The mature eggs that developed from the *dia-RNAi* egg chambers were significantly smaller than in controls (Fig. 3G). These data suggest that depletion of Dia alters ring canal structure and stability throughout oogenesis.

**Fig. 3:**
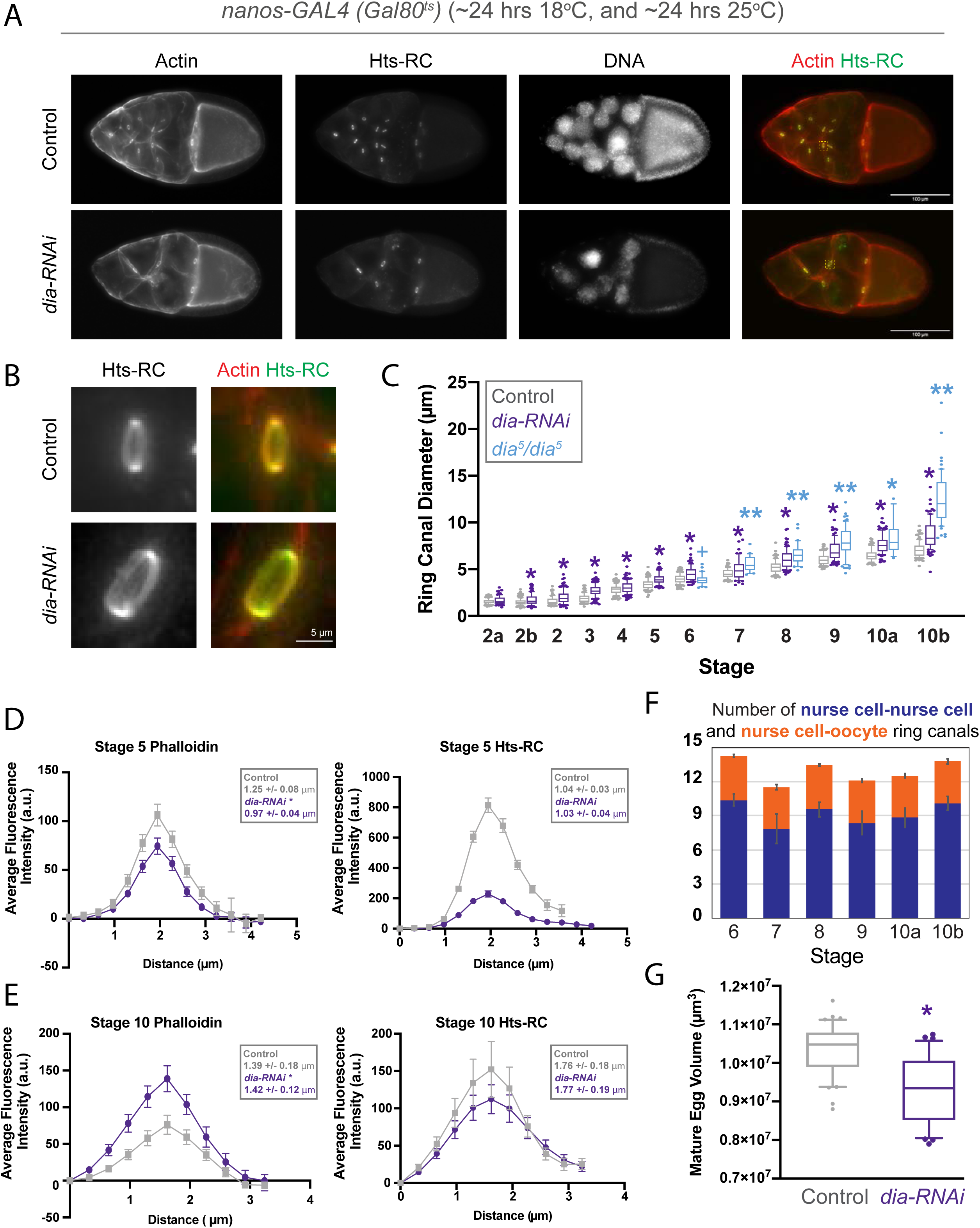
Depletion or mutation of the formin, Diaphanous, leads to nurse cell multinucleation and increased ring canal diameter. (A) Maximum intensity projections of control and *dia-RNAi* egg chambers. (B) Magnified view of the ring canals indicated by the boxed regions in A. (C) Box and whiskers plot showing the 10-90^th^ percent values for the diameter of ring canals connecting nurse cells in control, *dia-RNAi*, and *dia^5^/dia^5^* mutant germlines. Individual points represent values outside of that range. The DFS technique was used to generate *dia^5^/dia^5^* mutant germlines. This technique does not utilize a visible marker to identify mutant or wild type cells; instead, egg chambers containing nurse cells that have not undergone mitotic recombination will be heterozygous for the *ovo^D1^* mutation, which will cause them to degenerate around stage 5 of oogenesis. Therefore, only stage 6-10b egg chambers could be analyzed. n = 58-152 ring canals/stage for each condition. Asterisks indicate significant difference compared to control (p<0.05, K-S test); + indicates significant difference compared to *dia-RNAi* (p<0.05, K-S test), and double asterisk indicates significant different compared to both control and *dia-RNAi* (p<0.05, K-S test). (D,E) Average fluorescence intensity of phalloidin and Hts-RC staining in ring canals of (D) stage 5 and (E) stage 10 control and *dia-RNAi* egg chambers (n=49 for control, n=51 for *dia-RNAi* in D; n=12 for control, n=19 for *dia-RNAi* in E). Error bars are SEM. Average full width at half maximum +/− SEM is shown for each stain in each condition. (F) Average number of ring canals connecting nurse cells (blue) or nurse cells and the oocyte (orange) at each stage in *dia-RNAi* egg chambers. There should be a total of 15 ring canals per egg chamber at each stage. n=6-12 egg chambers/stage. Error bars are SEM. (G) Box and whiskers plot showing the 10-90^th^ percent values for the volumes of mature eggs. n=41 eggs for control and n=39 eggs for *dia-RNAi*. Asterisk indicates significant difference compared to control (p<0.0001, 2-tailed t-test). For all experiments, Dia was depleted using the nanos-GAL4 driver under the control of the temperature sensitive repressor *(GAL80^ts^); nanos-GAL4* shows a peak of expression within the germarium and again during mid-oogenesis.

In order to determine whether the reduced number of ring canals was due to defects in germ cell divisions within the germarium or to nurse cell fusion during later stages of oogenesis, the number of ring canals and nurse cell nuclei was monitored in *dia-RNAi* egg chambers. Normally, each egg chamber should contain 15 nurse cells (and one oocyte) and 15 ring canals. In the *dia-RNAi* egg chambers with fewer than 15 ring canals, some contained an equal number of nurse cell nuclei, which would suggest an early defect in germ cell division (such as failure of incomplete cytokinesis) during formation of the germ cell cyst (Fig. S1A). A similar phenotype has been observed in egg chambers with germline mutations in the myosin regulatory light chain (Wheatley et al., 1995). However, there were also many egg chambers that contained more nurse cell nuclei than ring canals (Fig. S1A), suggesting a later nurse cell fusion event. Therefore, depletion of Dia can lead to both defects in incomplete cytokinesis during germ cell division, and ring canal or membrane instability, sometimes leading to nurse cell fusion.

The average diameter of ring canals in *dia-RNAi* egg chambers was significantly larger than in controls at almost all stages analyzed (Fig. 3C). To confirm that this phenotype was specific to depletion of Dia, the dominant female sterile (DFS) technique (Chou et al., 1993; Chou and Perrimon, 1996, 1992) was used to generate egg chambers that contained homozygous mutant (*dia^5^ FRT40/dia^5^ FRT40*) germ cells. Although not a null allele, *dia^5^* has been shown to significantly reduce expression of Dia (Homem and Peifer, 2009, 2008). The ring canals in these mutant germ cells were indeed significantly larger than in controls at most stages analyzed and were even significantly larger than ring canals from *dia-RNAi* egg chambers (Fig. 3C).

The increase in ring canal diameter observed in the *dia-RNAi* and mutant egg chambers might not be due to a specific role for Dia in regulating ring canal size. Instead, it could be caused by reduced closure of the contractile ring during incomplete cytokinesis or because larger, multinucleate nurse cells produced by fusion may, by their nature, have larger ring canals. To uncouple these two aspects of the phenotype, we plotted the diameter of ring canals connecting nurse cells only in egg chambers that contained the appropriate number of ring canals. These ring canals were still significantly larger than controls at most stages analyzed (Fig. S1B), suggesting that Dia plays a role in limiting ring canal expansion after incomplete cytokinesis. However, it is also possible that more modest defects in incomplete cytokinesis, such as reduced contractile ring closure, generate an initially larger ring canal that then expands at a normal rate.

### Depletion of ArpC2 or Dia after germline cyst formation alters ring canal size

Even in the earliest stages of oogenesis, within the anterior structure of the ovariole, the germarium, egg chambers depleted of ArpC2 or Dia had significantly larger ring canals. These early size differences likely reflect a role for these two nucleators during incomplete cytokinesis of the cystoblast cells, consistent with recent work showing that a balance between the two nucleators must be maintained during cytokinesis (Chan et al., 2019). However, this early ring canal size difference makes it difficult to specifically assess the contribution of the Arp2/3 complex and Dia during ring canal expansion.

In order to isolate later stages of oogenesis, we used the *matαTub-GAL4* driver, whose expression begins around stage 2 of oogenesis (Fig. 1B; Hudson and Cooley, 2014), to reduce the levels of ArpC2 or Dia in the germline. The efficiency of mRNA knock-down was verified by qRT-PCR (Fig. S2A). Depletion of ArpC2 after formation of the germ cell cyst again led to significant variability in ring canals size. At stage 6 and 7, average ring canal diameter was significantly larger than in controls, whereas beginning at stage 9, average ring canal diameter was significantly reduced. Although not as frequent, lumenless ring canals were still observed (Fig. 4A,B). In contrast, depletion of Dia under the same conditions led to larger ring canals at most stages analyzed (Fig. 4A,B). Analysis of mature egg volume demonstrated that germline depletion of ArpC2 led to formation of a smaller mature egg, but depletion of Dia did not significantly affect final egg volume (Fig. 4C). These data indicate that both the Arp2/3 complex and Dia play a role in determining ring canal size after formation of the germ cell cyst, but that the ring canal enlargement observed in the *dia-RNAi* egg chambers does not significantly affect final egg volume.

**Figure 4:**
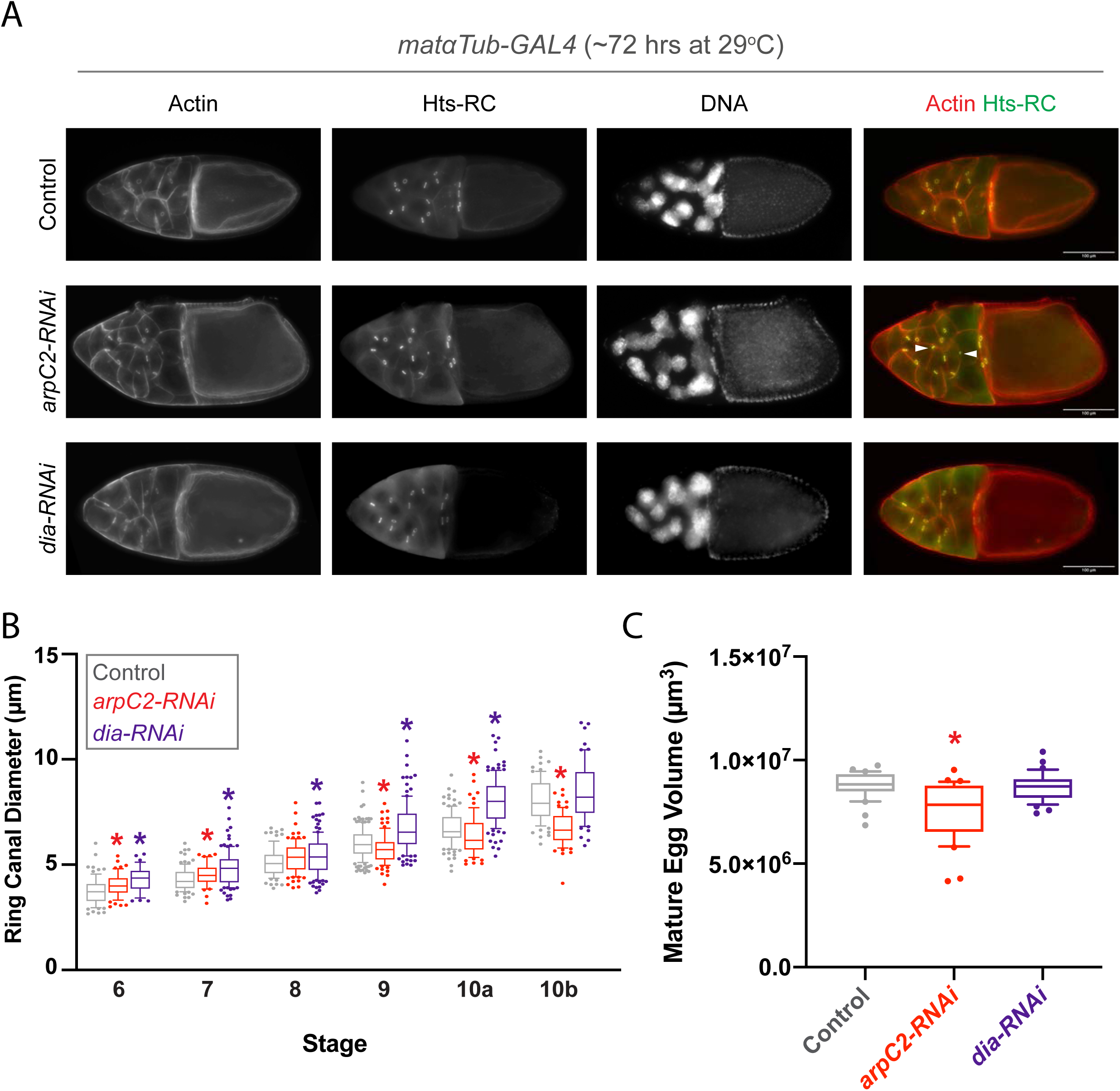
Both Diaphanous and the Arp2/3 complex are required to modulate ring canal size during oogenesis. (A) Maximum intensity projections of control, *arpC2-RNAi*, and *dia-RNAi* egg chambers. Arrowheads indicate small, lumenless ring canals. (B) Box and whiskers plot showing the 10-90^th^ percent values for the diameter of ring canals connecting nurse cells in control, *arpC2-RNAi*, and *dia-RNAi* egg chambers. Individual points represent values outside of that range. n = 33-122 ring canals/stage for each condition. Asterisks indicate significant difference compared to control (p<0.05, K-S test). (C) Box and whiskers plot showing the 10-90^th^ percent values for the volumes of mature eggs. n=30 eggs for control, n=35 eggs for *arpC2-RNAi*, and n=39 eggs for *dia-RNAi*. Asterisk indicates significant difference compared to control (p<0.0001, one-way ANOVA with Tukey’s post hoc). ArpC2 or Dia was depleted beginning at stage 2 of oogenesis using *matαTub-GAL4*.

### Expression of activated Dia leads to smaller and collapsed ring canals

The Arp2/3 complex and formin family members have been shown to function both cooperatively (Isogai et al., 2015) and antagonistically (Burke et al., 2014; Rotty et al., 2015; Suarez et al., 2015). If there is a balance or competition between the two actin nucleators, the phenotype produced by the depletion of one nucleator should be at least partially due to the over-activation of the other. Expression of a GFP-tagged, activated form of Dia (either GFP-Dia^ΔDAD^, which lacks the autoinhibitory domain, or GFP-Dia^FH3FH1FH2^, which contains point mutations in three of the essential regulatory domains; Homem and Peifer, 2009) can be used to upset the potential balance between Arp2/3 and Dia. Egg chambers expressing activated Dia beginning at stage 2 (using *matαTub-GAL4*) had a fairly normal number of nurse cell nuclei (Fig. 5A,B), but there was a progressive increase in the number of lumenless ring canals and a loss of nurse cell-nurse cell ring canals from stage 6-10b (Fig. 5A,C). By stage 10b, these egg chambers contained an average of only ~6-9 ring canals connecting nurse cells (instead of 11), and an average of 2-3 of these ring canals lacked a clear lumen (Fig. 5C). In addition, we observed abnormal actin structures or spikes on the surface of some of the nurse cells expressing activated Dia (Fig. 5A). These observations suggest that expression of activated Dia may disrupt the balance in nucleator activity and either block ring canal expansion or actively induce ring canal constriction to the degree that the ring canals become lumenless and are eventually lost over the course of oogenesis. These smaller, lumenless ring canals are reminiscent of those seen the *arpC2-RNAi* condition (Fig. 2), but the progressive ring canal loss suggests that this condition is more severe.

**Figure 5:**
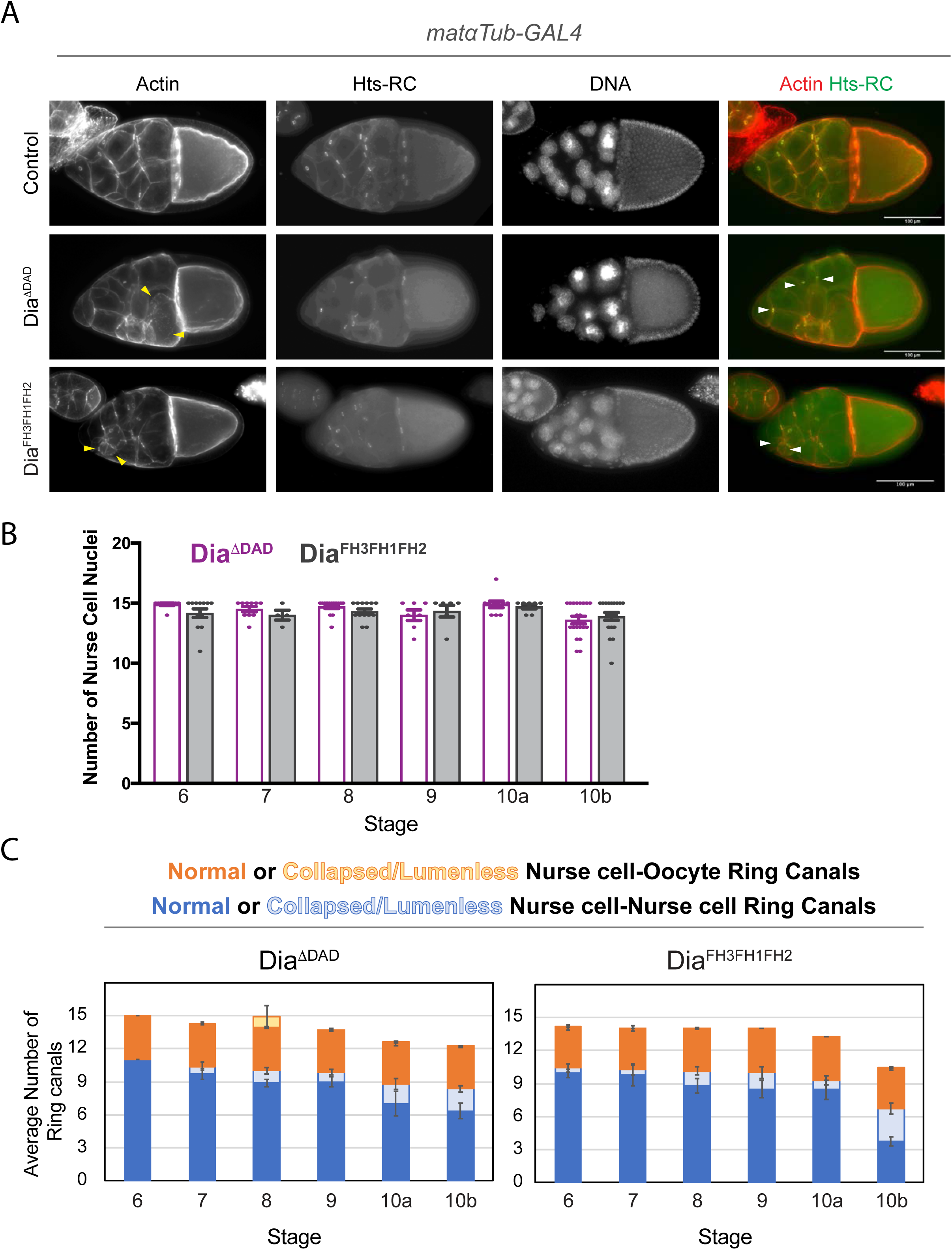
Expression of a constitutively active form of Diaphanous leads to ring canal collapse and and NC fusion. (A) Maximum intensity projections of control, Dia^ΔDAD^, and Dia^FH3FH1FH2^-expressing egg chambers. White arrowheads indicate small, lumenless ring canals; yellow arrowheads indicate abnormal actin structures. (B) Average number of nurse cell nuclei in Dia^ΔDAD^-or Dia^FH3FH1FH2^ – expressing egg chambers. (C) Average number of normal and collapsed/lumenless ring canals in egg chambers expressing Dia^ΔDAD^ or Dia^FH3FH1FH2^ in the germline. Error bars are SEM. Dia^ΔDAD^ and Dia^FH3FH1FH2^ were expressed beginning at stage 2 of oogenesis using *matαTub-GAL4*.

### Reducing Dia levels provides a modest rescue of the arpC2-RNAi phenotype

Over-expression of activated Dia demonstrated that altering the balance between these nucleators disrupts ring canal size and structure. If these nucleators are competing with or negatively regulating each other, then partially reducing the levels of one may be able to rescue the phenotype produced by depleting the other. During cytokinesis in the *C. elegans* embryo, partial depletion of the formin CYK-1 rescues depletion of the Arp2/3 subunit, ARX-2 (Chan et al., 2019). If the Arp2/3 complex and Dia are engaged in a similar form of competition in order to regulate ring canal size, then reducing the levels of Dia should rescue the *arpC2-RNAi* phenotype. We introduced a heterozygous mutation in Dia (*dia^5^ FRT40/+*) into the *arpC2-RNAi* condition (*dia^5^ FRT40/+; arpC2-RNAi)*, and performed a slightly weaker depletion of ArpC2 than was done previously (incubating the flies for ~48 hours at 25°C compared to ~72 hours at 29°C, which was shown in Fig. 2). Focusing on the later stages of oogenesis, just prior to nurse cell dumping, we again found that depletion of ArpC2 led to the formation of smaller (Fig. 6A,B) and lumenless (Fig. 6A,C) ring canals and the production of smaller mature eggs (Fig. 6D). In egg chambers heterozygous for the Dia mutation (*dia^5^ FRT40/+*), there was a modest yet significant reduction in ring canal diameter at stage 10b (Fig. 6B), but this did not result in a significant difference in mature egg volume compared to control (Fig. 6D). When Dia was reduced in the *arpC2-RNAi* background (*dia^5^ FRT40/+; arpC2-RNAi)*, average ring canal diameter was further reduced (but only significantly different at stage 10a), when compared to *arpC2-RNAi* alone (Fig. 6B), but there was a significant rescue of the number of lumeneless ring canals at stages 9 and 10a (Fig. 6C). The mature eggs produced did not differ in volume when compared to *arpC2-RNAi* alone (Fig. 6D). These data suggest that reducing Dia levels could provide a modest rescue of the *arpC2-RNAi* phenotype, but that both nucleators are essential to regulate ring canal size and stability in order to support cytoplasmic transfer during nurse cell dumping.

**Figure 6:**
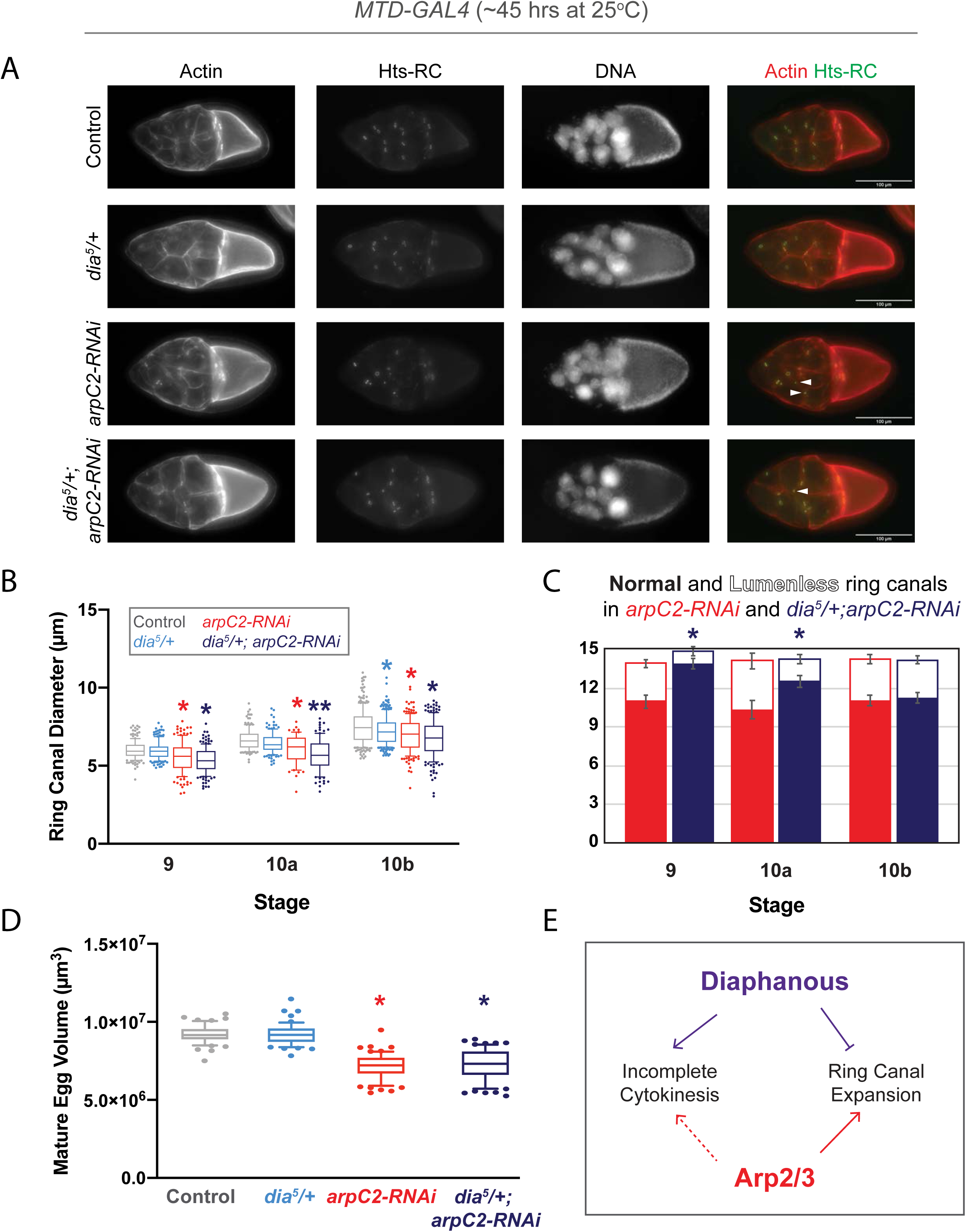
Reducing Dia levels provides a modest rescue of the *arpC2-RNAi* phenotype. (A) Maximum intensity projections of control, *dia^5^/+*, *arpC2-RNAi*, and *dia^5^/+; arpC2-RNAi* egg chambers. Arrowheads indicate small, lumenless ring canals. (B) Box and whiskers plot showing the 10-90^th^ percent values for the diameter of ring canals connecting nurse cells in control, *dia^5^/+*, *arpC2-RNAi*, and *dia^5^/+; arpC2-RNAi*. Individual points represent values outside of that range. n = 33-122 ring canals/stage for each condition. Asterisks indicate significant difference compared to control (p<0.05, K-S test). (C) Average number of normal and lumenless ring canals in *arpC2-RNAi* and *dia^5^/+; arpC2-RNAi* egg chambers in stages 9, 10a, and 10b. Error bars are SEM. Asterisk indicates significant difference in the number of lumenless ring canals when compared to *arpC2-RNAi* alone (p<0.05, 2-tailed t-test) (D) Box and whiskers plot showing the 10-90^th^ percent values for the volumes of mature eggs. n = 58 eggs for control, n=59 eggs for *dia^5^/+*, n = 69 eggs for *arpC2-RNAi*, and n = 63 eggs for *dia^5^/+; arpC2-RNAi*. Asterisk indicates significant difference compared to control (p<0.0001, one-way ANOVA with Tukey’s multiple comparison post hoc). All crosses were done using *MTD-GAL4*.

## Discussion

Here, we have demonstrated that altering the levels of two actin nucleators, the Arp2/3 complex, or the formin, Diaphanous (Dia), in the germline lead to distinct defects in oogenesis. Depletion of the Arp2/3 complex subunit, ArpC2, led to variation in ring canal size throughout oogenesis. During Phase I, ring canals were larger on average in the *arpC2-RNAi* egg chambers, but by Phase II, they become smaller than controls (Fig. 2,4). In contrast, depletion of Dia consistently caused over-expansion of the germline ring canals (Fig. 3,4); this over-expansion could be uncoupled from nurse cell fusion (Fig. S1A) and a role in incomplete cytokinesis (Fig. 4). By depleting ArpC2 or Dia after the germline cyst has formed, we confirmed a role for both actin nucleators in modulating ring canal size (Fig. 4). Constitutive activation of Dia led to a progressive phenotype in which ring canals collapsed and eventually disappeared (Fig. 5). Finally, because reduction of Dia could partially, but not fully, rescue the *arpC2-RNAi* phenotype (Fig. 6), the two nucleators likely cooperate to regulate ring canal size in the germline (Fig. 6E).

### The Arp2/3 complex and Diaphanous likely both play a role in regulating incomplete cytokinesis and ring canal expansion

Our data suggest that both the Arp2/3 complex and Dia may be necessary to regulate the efficiency of contractile ring closure during incomplete cytokinesis, as has been seen in other systems. In the *C. elegans* embryo, the formin, CYK-1, plays a direct role in nucleating actin filaments to form the contractile ring and promote early stages of constriction (Chan et al., 2019; Davies et al., 2014), whereas the Arp2/3 complex provides a more indirect contribution to the process by inhibiting excess formin (CYK-1) activity in the furrow region (Chan et al., 2019; Sun et al., 2011). In addition, both Formin-2 and the Arp2/3 complex have been implicated in regulating cytokinesis during asymmetric division of the mouse oocyte (Dumont et al., 2007; Sun et al., 2011). Although our data cannot uncouple a role during incomplete cytokinesis from one during the very earliest stages of ring canal size determination, it does suggest that cooperation between these types of actin nucleators could be commonly used in traditional, asymmetric, and incomplete cytokinesis.

Ring canal expansion is a unique process that, at first glance, may resemble cytokinesis in reverse, but upon closer inspection, it requires a distinct set of cellular machinery. The rates of actin turnover within the ring, at least at stage 10a, are within the range of those reported at the leading edge of the lamellipodia of migrating cells (Kelso et al., 2002; McGrath et al., 1998; Theriot and Mitchison, 1991); therefore, a more accurate model may be to compare ring canal expansion to migration of a continuous leading edge outward from the lumen, as has been proposed (Kelso et al., 2002). If viewed through this lens, the cooperation between the Arp2/3 complex and Dia is not surprising. Recent work has shown that formation of lamellipodia in EGF-treated HeLa cells depends on both the formin, mDia1, and the Arp2/3 complex (Isogai et al., 2015). Although the Arp2/3 complex is the primary actin nucleator that drives membrane protrusion in the context of cell migration (Pollard, 2007), it requires existing, and preferably newly-formed actin filaments for full activation (Blanchoin et al., 2000; Bugyi and Carlier, 2010; Ichetovkin et al., 2002). In this context, the formin, mDia1, provides a source of new actin filaments to which the Arp2/3 complex could bind (Isogai et al., 2015). We could envision a similar sequential, yet cooperative relationship between the Arp2/3 complex and Dia within the germline in which the primary force driving ring canal expansion depends on the Arp2/3 complex, but activation of the Arp2/3 complex depends on the presence of Dia-nucleated mother filaments. Interestingly, Dia has been shown to promote localization of the Arp2/3 activators, SCAR and WASp, in the context of myoblast fusion (Deng et al., 2015), so there could be additional mechanisms to coordinate the activity of these two nucleators in the germline.

In other cellular contexts, the Arp2/3 complex and formins have been shown to have a more antagonistic or competitive relationship. In fission yeast, the Arp2/3 complex and the formin, Cdc12, compete for access to monomeric actin; the balance between the levels of these two nucleators determines the abundance of the structures that they build. Profilin is an essential regulator of this competition, as only formins are able to utilize profilin-actin to nucleate or elongate existing actin filaments (Burke et al., 2014; Rotty et al., 2015; Suarez et al., 2015). Therefore, the levels of profilin in the cell can alter the balance between nucleators to maintain overall network homeostasis. Profilin (Chickadee) is essential for oogenesis; *chickadee* mutant egg chambers contain fewer than 15 nurse cells, most of which contain 2 or more nuclei, and display defects in the formation of the filopodia-like actin cables that restrict nuclear movement during nurse cell dumping (Cooley et al., 1992; Verheyen and Cooley, 1994). Additional studies will be required in order to determine how profilin levels affect the balance between Arp2/3 and Dia activity in the germline and further whether this balance is important during incomplete cytokinesis and/or ring canal expansion.

### The Arp2/3 complex and Dia could indirectly affect ring canal size and structure

Despite the strong ring canal phenotypes, we were surprised to find that depletion of ArpC2 or Dia did not lead to more dramatic changes in the levels of f-actin within the ring canals. One possibility is that the Arp2/3 complex and Dia cooperate to promote actin filament nucleation within the ring, but in addition to that role, these two modulate ring canal size more indirectly through regulation of cell-cell adhesion. Cadherin-based adhesion is essential for ring canal anchoring and nurse cell stability (Loyer et al., 2015; Oda et al., 1997; Peifer et al., 1993; White et al., 1998), and recent work suggests that phosphorylation of the E-cadherin binding protein, β-catenin, is required for ring canal expansion (Hamada-Kawaguchi et al., 2015). The Arp2/3 complex promotes formation of adhesion complexes and regulates their normal turnover (Bernadskaya et al., 2011; Kovacs et al., 2011, 2002; Tang and Brieher, 2012; Verma et al., 2012, 2004), and Dia, its homologs, and other members of the formin family of actin nucleators have been implicated in regulating junctional actin and adherens junction stability (Carramusa et al., 2007; Gauvin et al., 2015; Grikscheit et al., 2015; Grikscheit and Grosse, 2016; Homem and Peifer, 2008; Kobielak et al., 2004; Phng et al., 2015; Rao and Zaidel-Bar, 2016; Sahai and Marshall, 2002). Therefore, the Arp2/3 complex and Dia could be well-positioned to couple changes in the actin cytoskeleton with regulated cell-cell adhesion to modulate ring canal expansion. Further studies will be required in order to characterize junctional stability in the germline, the connection to ring canal structure and expansion, and the potential contribution of these nucleators to the process.

Dia has also been implicated in affecting myosin activity, which may provide an additional mechanism to regulate ring canal size and stability. Although myosin-based contractility has not been shown to be directly required for ring canal expansion, mutation in the myosin phosphatase (dMYPT), significantly reduces ring canal size (Ferreira et al., 2014; Ong et al., 2010; Wheatley et al., 1995; Yamamoto et al., 2013), suggesting that myosin activity might need to be actively suppressed to facilitate expansion. In the fly embryo, reducing the levels of Dia and Rho1 led to reduced myosin activity and increased protrusiveness, whereas expression of an activated form of Dia increased myosin activity and reduced protrusions (Homem and Peifer, 2008). In our hands, expression of an activated form of Dia produced egg chambers with abnormal, lumenless, or missing ring canals. This phenotype could be due to defects in the balance between Arp2/3 and Dia actin activity; alternatively, the small ring canals could be due to excessive myosin activity. Genetic interaction and localization experiments will be necessary in order to determine whether Dia could be upstream of myosin phosphorylation in the germline and also determine the connection between myosin activity, adherens junction stability, and ring canal expansion.

The coordinated activity of multiple actin-binding and bundling proteins is also likely required to stabilize ring canals and promote their expansion. A number of actin-binding proteins that contribute to ring canal formation, stability, or expansion have been identified, including Filamin/Cheerio, Hts-RC, and Kelch (Kelso et al., 2002; Robinson et al., 1997, 1994; Robinson and Cooley, 1997), and some of these actin-binding proteins may even impact the activity of the nucleators. For example, during dorsal closure in the developing fly embryo, the balance between the level of the actin-binding protein, Enabled (Ena), and Dia determines whether a lamellipodia or filopodial structure will be formed in the leading edge cell (Homem and Peifer, 2009). Ena has been reported to localize near the germline ring canals (Gates et al., 2009), which would make it a potential candidate to modulate Dia activity during oogenesis, as well as to directly impact actin dynamics.

### The Arp2/3 complex and Dia may regulate other germline actin structures

Although our work has focused on the role of these two nucleators in regulation of the germline ring canals, there are other essential actin-based structures that promote efficient cytoplasmic transfer during oogenesis. For example, during stage 10b, a set of filopodia-like actin cables assembles from the basolateral surfaces of the nurse cells. These actin cables restrict the movement of the nurse cell nuclei during dumping such that they do not obstruct the ring canal and block transport (Guild et al., 1997; Gutzeit, 1986; Huelsmann et al., 2013). Mutation of Arp2/3 complex members did not disrupt formation of the actin cables (Hudson and Cooley, 2002), but to our knowledge, the role for Dia in cable formation or extension has not been tested. Further, it is possible that depletion of Arp2/3 or Dia could disrupt cortical actin structure, thereby indirectly affecting actomyosin based contractility, which is the primary driving force for nurse cell dumping (Ferreira et al., 2014; Gutzeit, 1986; Spradling, 1993; Wheatley et al., 1995).

Our data have characterized the role for two conserved actin nucleators in regulation of the size, stability, and expansion of germline intercellular bridges. Future studies are required to determine how these nucleators are regulated as well as the specific mechanisms by which they modulate the size and stability of these essential cellular connections.

## Supplemental Figure Legends

**Figure S1: Depletion or germline mutation of Diaphanous leads to ring canal expansion independent of defects in cytokinesis or nurse cell fusion.** (A) Number of ring canals and nurse cell nuclei in individual *dia-RNAi* egg chambers. A line with a 1x slope is included to show that in normal egg chambers, there should be a 1:1 correlation between ring canal number and the number of nurse cell nuclei. (B) Box and whiskers plot showing the 10-90^th^ percent values for the diameter of ring canals connecting nurse cells in control, *dia-RNAi*, and *dia^5^/dia^5^* mutant germlines (these data represent a subset of those shown in Fig. 3C); only the measurements from egg chambers containing 15 ring canals are included. Individual points represent values outside of that range. n = 22-120 ring canals/stage for each condition. Asterisks indicate significant difference compared to control (p<0.05, K-S test).

**Figure S2: UAS-RNAi transgenes effectively reduce ArpC2 or Dia mRNA.** (A) Average level of *arpC2* or *dia* mRNA remaining in whole ovary extract assessed by qRT-PCR (n=3). Error bars indicate SEM. ArpC2 or Dia was depleted beginning at stage 2 of oogenesis using *matαTub-GAL4*.

## Materials and Methods

### Fly Genetics and Maintenance

For full genotypes and maintenance conditions, see Table S1. The following stocks were obtained from the Blooming *Drosophila* Stock Center (BDSC): the maternal triple driver (*MTD-GAL4*) (*otu-GAL4; nos-GAL4; nos-GAL4*; #31777), *MatαTub-GAL4* (#7062 or #7063), *UAS-dia-RNAi* (#35479), *UAS-arpC2-RNAi* (#43132), *dia^5^ FRT40A/Cyo* (#9138), *ovoD1 FRT40A* (#2121), *UASp-dia.DeltaDad.EGFP* (#56752), and, *UASp-dia.FH3FH1FH2.EGFP* (#56753). The following lines were generated through genetic crossing: GAL80^ts^; nos-GAL4 (Kline et al., 2018), *hs-FLP; dia^5^ FRT40/Cyo*, *dia^5^ FRT40/Cyo; arpC2-RNAi/TM6c*. To induce *dia^5^* mutant germline clones, L2/L3 stage larvae were heat shocked at 37°C for 2 hours followed by a 22 hour recovery at 25°C. This procedure was then repeated one additional time before returning flies to the 25°C incubator.

### Quantification of RNA Levels

Relative expression of Dia and ArpC2 RNA was quantified as described previously (Kline et al., 2018). Briefly, total RNA was extracted from ovaries of well-fed female flies using the RNeasy Plus Mini kit (Qiagen). The QuantiNova SYBR Green RT-PCR kit (Qiagen) was used to reverse transcribe cDNA, and the following primers were used: ArpC2 – 5’ – CTTCGACGGCGTCCTTTATCA – 3’ and 5’ – GCCATATTCACGCTTCAGCAAC – 3’; Dia – 5’ – GGCTCCTGGACAGTCTGTTC – 3’ and 5’ – GGTTCCTCCCTTGGACTTCG – 3’ (FlyPrimerBank(Hu et al., 2013)). GAPDH was used as the control: 5’ – AAATCGCGGAGCCAAGTAGT-3’ and 5’ – CACGATTTTCGCTATGGCCG – 3’ (Giuliani et al., 2014). Quantification of Dia and ArpC2 mRNA levels was calculated using the using the 2-ΔCq method (Kline et al., 2018; Livak and Schmittgen, 2001). Data in Fig. S2A represent the average of 3 independent experiments; each qRT-PCR reaction was done in triplicate for each experiment.

### Ovary Dissection and Staining Procedures

Female flies of the indicated genotypes were incubated with fresh ground yeast in the presence of males at 18°-29°C for 24-72 hours as indicated in Table S1. After incubation, ovaries were dissected in Schneider’s media (Genesee Scientific) and fixed in 4% EM grade formaldehyde (Polysciences), which was diluted in either PBS or PBS + 0.1% Triton X-100 (PBT) for 15 minutes. Tissue was washed in PBT and PBT3 (PBS+0.3% Triton X-100) and stained with DAPI (1:500, 1 mg/mL stock, D3571 ThermoFisher Scientific), TRITC- or FITC-conjugated phalloidin (1:500, ECM Biosciences), and/or an antibody targeting Hts-RC diluted in PBT3 + 5 mg/mL BSA. The HtsRC antibody (1:20, hts RC) was obtained from the Developmental Studies Hybridoma Bank (DSHB). Secondary antibodies were conjugated to Alexa Fluor 488/DyLight 488 or Alexa Fluor 555 (donkey anti-mouse or goat anti-mouse from Invitrogen or Jackson ImmunoResearch); they were diluted 1:100-1:200 in PBT3 + 5mg/mL BSA. SlowFade Antifade (ThermoFisher Scientific) was used to mount tissue on slides.

### Mature Egg Measurements

Females were incubated with fresh ground yeast in the presence of males for ~24 hours prior to moving them to apple juice plates with wet yeast. Mature eggs were collected at the indicated times and temperatures (Table S1), rinsed from apple juice plates with distilled water, and imaged using the 10x objective on a standard Zeiss compound microscope equipped with a ProgRes MF camera (Jenoptik). Images were captured using the ProgRes Capture Pro software (Jenoptik). Fiji/ImageJ was utilized to measure the length and width of the mature eggs, and the following formula was used to calculate the volume of the mature eggs: Volume = 1/6(π)(width)^2^(length) as done previously (Kline et al., 2018).

### Imaging and Analysis

Most of the images were collected using a Leica DM5500B fluorescence microscope with DFC7000T camera controlled by Leica Application Suite X (LASX) software. z-stacks were collected through the thickness of the egg chambers (using the system optimized z-step separation). To quantify ring canal diameter from the Hts-RC stain, Fiji/ImageJ was used to locate the z-plane where each ring canal was in focus, and the line tool was used to measure the outer diameter of each ring canal. Ring canals lacking a clear lumen were not included in the outer diameter measurements. Linescan analysis was performed as described previously (Kline et al., 2018). Statistical analyses (two-tailed t-test, one-way ANOVA with Tukey’s post-hoc analysis, or Kolmogorov-Smirnov test) were performed using the Prism 8 software (GraphPad).

## Acknowledgements

We would like to thank Kayla Harpold for assistance with data collection and Kristin Sherrard for helpful suggestions on the manuscript. This work was supported by the National Institutes of Health (NIH R15HD084243-01A1). Stocks obtained from the Bloomington *Drosophila* Stock Center (NIH P40OD018537) were used in this study. The hts RC antibody developed by L. Cooley was obtained from the Developmental Studies Hybridoma Bank, created by the NICHD of the NIH and maintained at The University of Iowa, Department of Biology, Iowa City, IA 52242.

