## Supplementary material for "The Arp2/3 complex and the formin, Diaphanous, are both required to regulate the size of germline ring canals in the developing egg chamber": Table S1

| Genotype | Temperature cross raised (°C) | Incubation temperature prior to dissection (°C) | Incubation time prior to dissection (hours) | Antibody | Associated Figures |
|---|---|---|---|---|---|
| *otu-GAL4/w[1118]; nos-GAL4/+; nos-GAL4/+* | 25 | 29 | 72 | Hts-RC (1:20) | Fig. 2A-F |
| *otu-GAL4/+; nos-GAL4/+; nos-GAL4/UAS-arpC2-RNAi* | 25 | 29 | 72 | Hts-RC (1:20) | Fig. 2A-F |
| *otu-GAL4/w[1118]; nos-GAL4/+; nos-GAL4/+* | 25 | 29 | 65.5 | | 2G |
| *otu-GAL4/+; nos-GAL4/+; nos-GAL4/UAS-arpC2-RNAi* | 25 | 29 | 65.5 | | 2G |
| *w[1118]/+;Gal80ts/+; nanos-GAL4/+* | 18 | 18, 25 | 24, 24 | Hts-RC (1:20) | Fig. 3A-F |
| *Gal80ts/UAS-dia-RNAi; nanos-GAL4/+* | 18 | 18, 25 | 24, 24 | Hts-RC (1:20) | Fig. 3A-F |
| *hsFLP/+;dia[5] FRT40/dia[5] FRT40* | 25 | 25 | 44-72 | Hts-RC (1:20) | Fig. 3A-F |
| *w[1118]/+;Gal80ts/+; nanos-GAL4/+* | 18 | 25 | 50 | | 3G |
| *Gal80ts/UAS-dia-RNAi; nanos-GAL4/+* | 18 | 25 | 50 | | 3G |
| *w[1118]/+;;matαTub-GAL4/+* | 25 | 29 | 72 | Hts-RC (1:20) | 4A-B |
| *matαTub-GAL4/UAS-arpC2-RNAi* | 25 | 29 | 72 | Hts-RC (1:20) | 4A-B |
| *UAS-dia-RNAi/+;matαTub-GAL4/+* | 25 | 29 | 72 | Hts-RC (1:20) | 4A-B |
| *w[1118]/+;;matαTub-GAL4/+* | 25 | 29 | 65.5 | | 4C |
| *matαTub-GAL4/UAS-arpC2-RNAi* | 25 | 29 | 65.5 | | 4C |
| *UAS-dia-RNAi/+;matαTub-GAL4/+* | 25 | 29 | 65.5 | | 4C |
| *w[1118]/+;matαTub-GAL4/+* | 25 | 29 | 43 | Hts-RC (1:20) | 5A |
| *matαTub-GAL4/+; UASp-dia.DeltaDad.EGFP/+* | 25 | 29 | 43 | Hts-RC (1:20) | 5A |
| *UASp-dia.FH3FH1FH2.EGFP/matαTub-GAL4* | 25 | 29 | 43 | Hts-RC (1:20) | 5A |
| *w[1118]/+;matαTub-GAL4/+* | 25 | 25 | 72 | Hts-RC (1:20) | 5B,C |
| *matαTub-GAL4/+; UASp-dia.DeltaDad.EGFP/+* | 25 | 25 | 72 | Hts-RC (1:20) | 5B,C |
| *UASp-dia.FH3FH1FH2.EGFP/matαTub-GAL4* | 25 | 25 | 72 | Hts-RC (1:20) | 5B,C |
| *otu-GAL4/w[1118]; nos-GAL4/+; nos-GAL4/+* | 25 | 25 | 45 | Hts-RC (1:20) | 6A-C |
| *otu-GAL4/+; nos-GAL4/dia[5]; nos-GAL4/+* | 25 | 25 | 45 | Hts-RC (1:20) | 6A-C |
| *otu-GAL4/+; nos-GAL4/+; nos-GAL4/UAS-arpC2-RNAi* | 25 | 25 | 45 | Hts-RC (1:20) | 6A-C |
| *otu-GAL4/+; nos-GAL4/dia[5]; nos-GAL4/UAS-arpC2-RNAi* | 25 | 25 | 45 | Hts-RC (1:20) | 6A-C |
| *otu-GAL4/w[1118]; nos-GAL4/+; nos-GAL4/+* | 25 | 25 | 47 | | 6D |
| *otu-GAL4/+; nos-GAL4/dia[5]; nos-GAL4/+* | 25 | 25 | 47 | | 6D |
| *otu-GAL4/+; nos-GAL4/+; nos-GAL4/UAS-arpC2-RNAi* | 25 | 25 | 47 | | 6D |
| *otu-GAL4/+; nos-GAL4/dia[5]; nos-GAL4/UAS-arpC2-RNAi* | 25 | 25 | 47 | | 6D |
| *w[1118]/+;Gal80ts/+; nanos-GAL4/+* | 18 | 18, 25 | 24, 24 | Hts-RC (1:20) | Fig. S1B |
| *Gal80ts/UAS-dia-RNAi; nanos-GAL4/+* | 18 | 18, 25 | 24, 24 | Hts-RC (1:20) | Fig. S1A,B |
| *hsFLP/+;dia[5] FRT40/dia[5] FRT40* | 25 | 25 | 44-72 | Hts-RC (1:20) | Fig. S1B |
| *w[1118]/+;;matαTub-GAL4/+* | 25 | 29 | 72 | | Fig. S2A |
| *matαTub-GAL4/UAS-arpC2-RNAi* | 25 | 29 | 72 | | Fig. S2A |
| *UAS-dia-RNAi/+;matαTub-GAL4/+* | 25 | 29 | 72 | | Fig. S2A |
