## Supplementary figures and images for "The Arp2/3 complex and the formin, Diaphanous, are both required to regulate the size of germline ring canals in the developing egg chamber"

### Figure S1

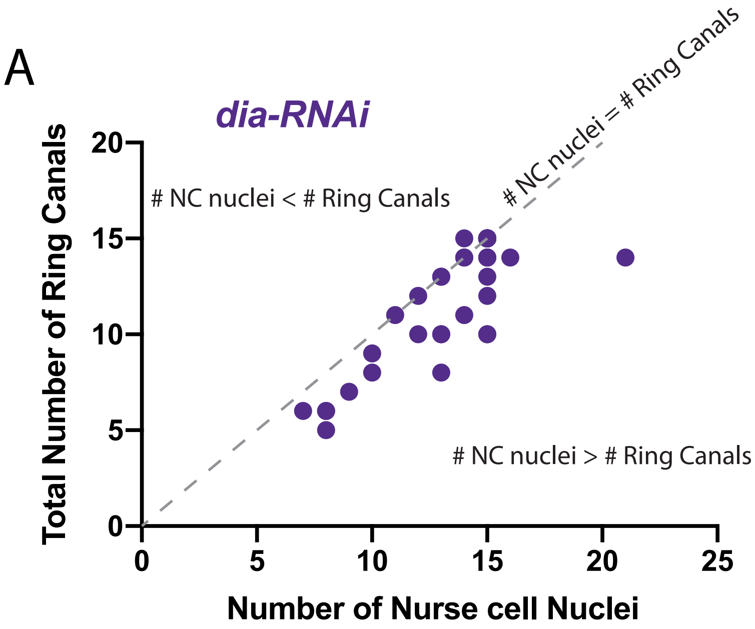

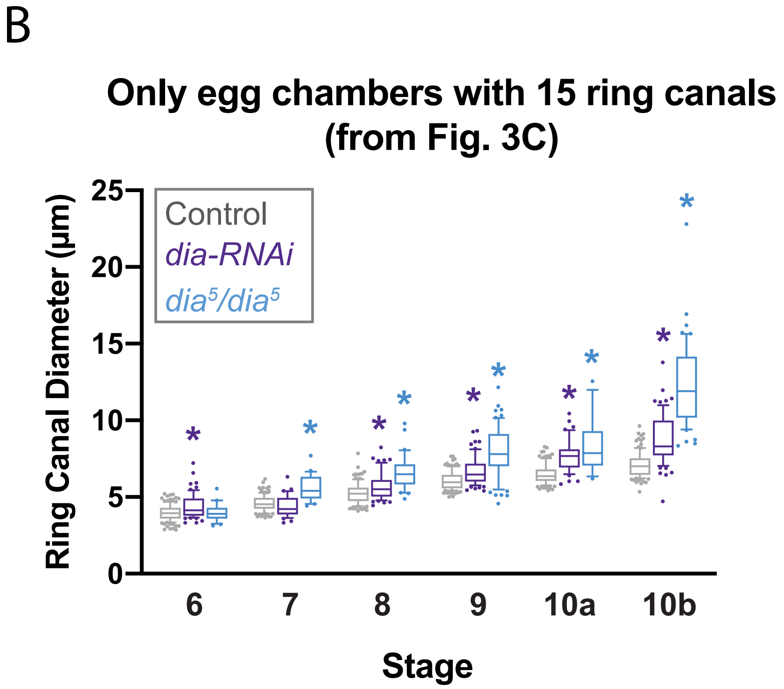

### Figure S2

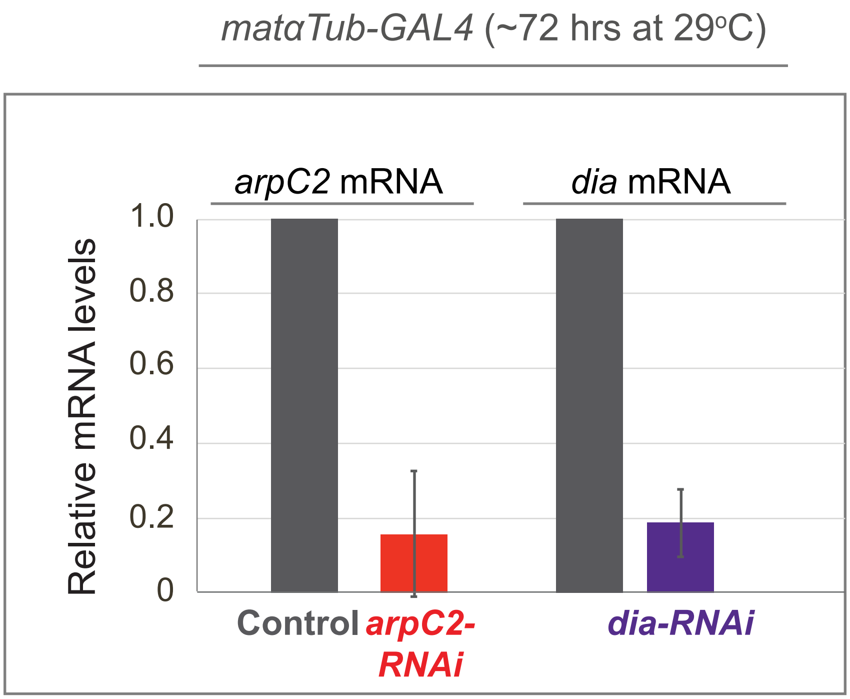
